# Mental Workload Estimation Using Wireless EEG Signals

**DOI:** 10.1101/755033

**Authors:** Quadri Adewale, George Panoutsos

## Abstract

Previous studies have shown that electroencephalogram (EEG) can be used in estimating mental workload. However, developing fast and reliable models for cross-task, cross-subject and cross-session classifications of workload remains a challenge. In this study, a wireless Emotiv EPOC headset was used to evaluate workload in two different mental tasks: n-back task and mental arithmetic task. 0-back task and 2-back task were employed as low and high workload in the n-back task while 1-digit and 3-digit addition were used as the two different workload levels in the arithmetic task. Using power spectral density as features, a fast signal processing and feature extraction framework was developed to facilitate real-time estimation of workload. Within-session accuracies of 98.5% and 95.5% were achieved in the n-back and arithmetic tasks respectively. Adaptive subspace feature matching (ASFM) was applied for cross-session, cross-task and cross-subject classifications. The feature adaptation provided average cross-session accuracies of 80.5% and 74.4% in the n-back and the arithmetic tasks respectively. An average cross-task accuracy of 68.6% was achieved while cross-subject accuracies were 74.4% and 64.1% in the n-back and arithmetic tasks respectively. The framework generalised well across subjects and tasks, and it provided a promising approach towards developing subject and task-independent models. This study also shows that a consumer-level wireless EEG headset can be applied in cognitive monitoring for real-time estimation of workload in practice.

## I. INTRODUCTION

Brain-computer interface (BCI) is mainly applied to aid disabled persons by using the brain signals for communication and control while bypassing auxiliary muscles or nerves [1]. However, the application of BCI for healthy patients is becoming increasingly popular and this is giving rise to a myriad of applications [2]. The application of BCI to obtain information about a user’s state by using arbitrary brain events without intending any voluntary control is called Passive BCI [2]. Passive BCI can be used to obtain information about a driver’s level of workload, stress or attentiveness. Consequently, the driver’s performance can be improved, and potential costly errors can be forestalled [3]. Furthermore, BCI has shown potential usefulness in avoiding accidents in industries by monitoring the mental state of workers [4]. In security surveillance, the level of attentiveness and concentration can also be monitored to ensure continuous safety [5]. Estimating workload in adaptive systems can generally facilitate task sharing and load shedding between human and machine to reduce operational errors.

Although there is no universal definition of workload [6], [2], mental workload can be determined by considering the task or the operator (human). The task-oriented approach considers the task characteristics and the condition of task performance in estimating workload. The human-oriented approach evaluates workload through the effect of the task performance on the human [7]. Psychologists are however more inclined to the latter approach – they view workload as the result of the interaction between work demands and human capacity [8]. As the workload increases, the task demand approaches the upper limit of human ability.

One way to measure workload is the use of performance measures. This method assumes that operator’s performance degrades with task demand. Performance degradation is evidenced in slower work pace and increased errors in the task. However, it could be costly to wait till performance degrades, especially in some critical tasks [9]. Nevertheless, performance measures are easy to justify and can be useful in building predictive models for operator functional state [6]. Subjective measures also provide a way of estimating workload. The operator gives a personal evaluation of the workload by filling a rating scale. Common rating scales include NASA Task Load Index (NASA-TLX) [7], Subjective Workload Assessment Technique (SWAT) [10], Rating Scale Mental Effort (RSME) [11] and Thurstonian Scale [12]. Rating scales can be used in many types of workload [7], they are easily prepared [12] and provide direct evaluation [10]. However, the requirement for an operator to repeatedly fill the scale may impose extra burden on him. In a bid to avoid such intermittent intrusion, an operator may fill the scale at a much later time, but this can lead to bias. The scale is also prone to falsification and faulty judgement [7]. Physiological correlates of workload have also been established in many literatures. Some of these measures include heart rate [13], blood pressure [14], chemical measures including sodium and potassium in saliva, cortisol in blood [14], heart rate variability [15], eye blink frequency [16], saccades [16], Electromyogram (EMG) [17] and EEG. Even though there seem not to be a best physiological indicator of workload, some studies showed EEG to be more promising compared to other indicators [18], [19].

Previous studies have shown correlation between workload and power in some EEG frequency bands. EEG frequency bands are commonly divided into delta (1-4Hz), theta (4-8Hz), alpha (8-12Hz), beta (12-25Hz) and gamma (>25Hz) [20]. Increase in workload was observed to cause reduction of power in alpha band, especially at the parietal regions [21]. Also, theta power was reported to increase with workload mostly in frontal regions [21]. Gamma [22], [23], delta and beta powers [13] increase with workload at the parietal and temporal regions.

In mental workload classification, a variety of machine learning methods have been employed. Some of these methods include support vector machine (SVM) [24], artificial neural network (ANN) [25] and hierarchical Bayes model [26]. SVM finds more application due to its ability to better generalise well and handle high-dimensional data [27]. However, due to the nonstationary nature of EEG signals, the performance of algorithms degrades when the training and test data are taken from different sessions and subjects. Hence the algorithm needs to be trained or adapted for every user and session. Some feature adaptation techniques have been proposed to control this degradation, but they are often computationally expensive and complex for real-time estimation of workload. Moreover, developing a model for cross-task classification remains a challenge. Furthermore, many of the previous studies on mental workload estimation employed large number of electrodes for recording the EEG signals, therefore reducing the comfortability of using EEG headsets in practical and online scenarios. Only a limited number of studies have recently investigated the feasibility of using wireless headsets with small number of electrodes for mental workload estimation [24]. To address these issues, we employed a wireless consumer-level Emotiv EPOC EEG headset with small number of electrodes to estimate mental workload. A simple signal processing and feature extraction technique was developed to facilitate practical and real-time application. The model was tested across eight (8) subjects in two different types of task – n-back task and arithmetic task. In addition, a fast domain adaptation technique called Adaptive Subspace Feature Machine (ASFM) [28] was applied to improve the model performance in cross-session, cross-task and cross-subject classifications. We compared the results from ASFM with those of SVM. Some subjective and performance indices of mental workload were also used to verify that the experimental design reflects different levels of workload

## II. METHOD

### A. Subjects

Eight (8) subjects (6 males and 2 females) participated in the EEG experiment at the Physiological Signal Processing Laboratory, Department of Automatic Control and Systems Engineering, University of Sheffield. Some of the participants were recruited through the University’s student volunteer list while others were friends of the research student. Participants were aged between 19 and 30 (M = 25 years; SD=3 years); where M = mean and SD = standard deviation. All subjects were right-handed, reported normal or corrected-to-normal vision, and had no history of any fatigue-related disorder. The experiment was performed in accordance with the University’s ethics guidelines, and participants were given written informed consent.

### B. Material

The EEG was recorded using a wireless Emotiv EPOC neuroheadset [29]. The Emotiv EPOC headset uses 14 electrodes with two additional electrodes for referencing (DRL) and noise cancellation (CMS) as shown in Fig. 1. All the available electrodes were used in this study. Saline liquid was applied to the electrodes before every experiment to keep the electrode impedances low to an acceptable level.

**Fig. 1.**
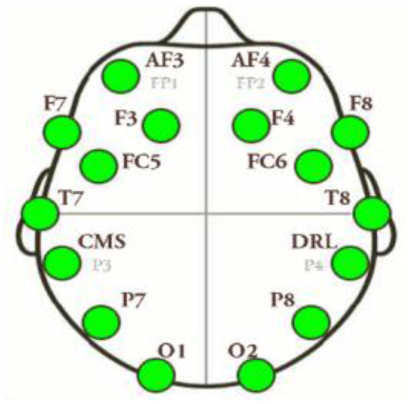
Electrode configuration of Emotiv EPOC [30]

The stimuli were presented on a monitor at a distance from participants’ eyes. Participants press keyboard to indicate if the letters or numbers displayed are targets or non-targets. They press ‘H’ (hit) if a target is displayed and ‘F’ (fail) for non-target. RSME scale [11] was also employed as a subjective measure of mental workload. RSME was chosen due to its higher sensitivity compared to other commonly employed methods like NASA-TLX [31]. Participants were asked to rate each task based on the perceived expended effort in solving that task. The scale ranges from 0 to 150. 0 depicts the lowest effort and 150 indicates the peak effort.

### C. Task

Two tasks were employed in the experiment and each task had two difficulty levels. The first task was n-back task and the second task was mental arithmetic task. N-back task has been extensively employed in cognitive workload studies because it can be used to vary workload by changing the number of letters (‘n’) a user should memorise. Moreover, changing the value of n does not change the rate of visual input or motor-cortex demands [7]. This is important to ensure that workload is being measured and not mere reaction to varying stimuli [32]. 0-back and 2-back tasks were used to represent low and high workloads respectively. In 0-back condition (low workload level), the target letter is ‘X’. Hence, the participant only needs to memorise the letter ‘X’ and does not need to update his memory as the task proceeds. For 2-back condition (high workload level), participant decides if the letter displayed currently is same as the letter displayed two sequences earlier. Hence, the participant updates his memory by memorising two previous letters as the sequence progresses. In both task levels, the participant presses the appropriate key to indicate if the letter is a target or not while paying attention to speed.

Mental arithmetic task has also been used to study cognitive workload [33] and is shown to have relationship with working memory [24]. The task requires a participant to perform arithmetic operations without any aid such as pen and paper or calculator. The answer from every arithmetic operation is stored in the participant’s brain and retrieved after some seconds when an answer is displayed. If the number displayed is the correct result from the last arithmetic operation, then such number is a target, else it is non-target. Two versions of arithmetic task were used. The first was 1-digit addition for low workload level and the second was 3-digit addition for high workload level. To regulate the effect of carry, a carry was always involved in both workload levels; e.g. (7 + 8) for 1-digit task, and (234 + 356) for 3-digit task.

### D. Stimuli

The stimuli were presented in Arial font style, white and size 50pt. The stimuli were displayed on a black background. Upper case letters were employed in the n-back tasks. Similar to the method used in [32], all letters were selected from consonants only and randomly presented. Vowel letters were not included to preclude participants from grouping the letters into sensible word or pattern, which may ease the workload. Each letter was displayed for 500ms, followed by a fixation cross for 2000ms (inter-stimulus interval) before the next letter. For both levels of n-back task, 120 trials were used and 30% of the trials were targets. The n-back task design is illustrated in Fig. 2. ‘T’ denotes ‘Target’, ‘NT’ denotes ‘None-Target’ and ‘s’ represents time in seconds.

**Fig. 2.**
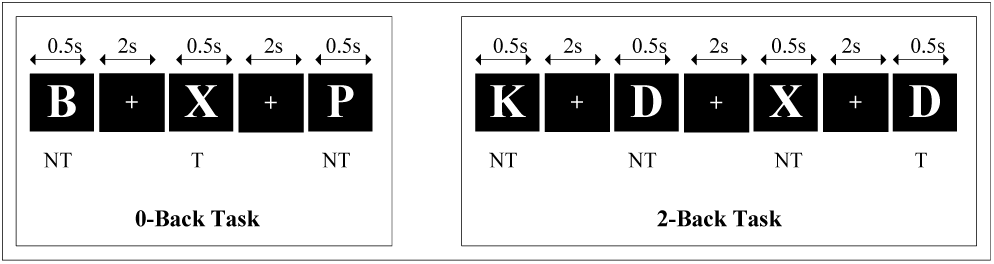
N-back tasks

For the mental arithmetic tasks, the two numbers to be added (trial) were displayed together for 5000ms followed by a fixation cross of 4000ms. Another single number (answer) was then presented for 2000ms and the participant indicates whether the number is a correct result of the last trial or not. A fixation cross was displayed for 2000ms before the next trial. 25 trials were used for both task levels and 44% of the trials were targets. Fig. 3 shows the 3-digit arithmetic task.

**Fig. 3.**
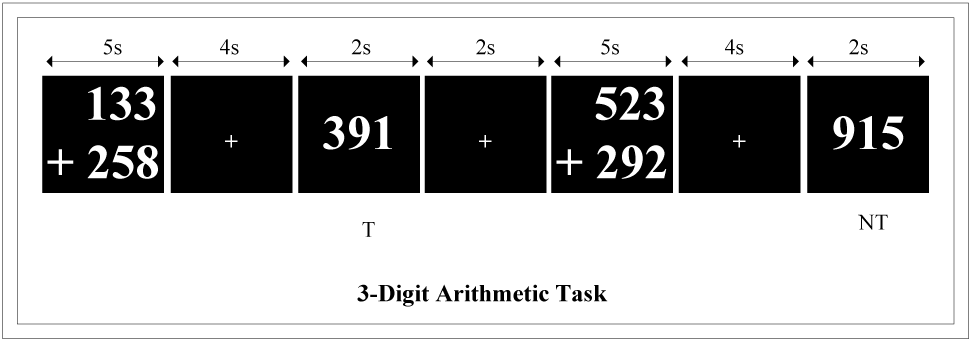
Arithmetic tasks

### E. Design

The experiment was designed to allow evaluation of model performance across different tasks. Each subtask (0-back, 2-back, 1-digit arithmetic and 3-digit arithmetic) was presented in a 5-minute block. Hence, a session of the experiment was comprised of four blocks. Markers were sent to the EEG recording panel to mark the beginning and end of each block. After each block of task, the participant could have a break before moving on to the next block. Before each session, there was a baseline block where the subject quietly fixated on a cross for 30 seconds. The response times and accuracies of answers were also recorded as dependent measures. The RMSE scale was filled by the participant after each block to subjectively rate the mental workload for each subtask.

Three of the eight participants were asked to repeat the tasks after one week; the tasks were presented in a counterbalanced order. The repetition was to enable the evaluation of robustness of the model to the nonstationarity of EEG. The whole experiment was designed and implemented in MATLAB with the aid of Cogent 2000 [34].

### F. Procedure

As the subjects arrived for the experiment, they were made to fill the screening form to ascertain if they were eligible for the experiment. After the screening, participant went on to sign informed consent form. Then, the tasks involved in the experiment were clearly explained to the subject. The EEG headset was worn on the participant’s head and proper connection of the electrodes to the scalp was ensured by observing the Emotiv panel throughout the experiment.

Before the actual task, participants practised 2-back task and 3-digit mental arithmetic task several times until they were confident with the task. Participant was told to pay attention to speed and accuracy during the practice and main experiment. The participant was able to see his performance after every practice. Thereafter, the participant moved on to do the main tasks. The instructions about the tasks were displayed on the monitor at necessary stages in the experiment. First, the participant fixated on a cross on the screen for 30 seconds without any movement or much eye blinking. Then, the participant moved on to perform a 5-minute block of the 0-back task. After the first block of task, participant could take a break before moving to the next task. Also, the participant was required to rate the 0-back task on the RMSE scale before proceeding. Participant then moved on to the second, third and fourth blocks to solve the 2-back, 1-digit arithmetic and 3-digit arithmetic tasks respectively. The participant took break after every block and rated the workloads on RMSE scale.

### G. Data Analysis

#### 1) Data Processing

The EEG data were sampled at 128Hz. All the 14 channels were used and referenced absolutely to the left mastoid. The raw EEG measurement in edf format was imported into MATLAB and converted to structure array (struct). The data from the fourteen channels were accessed by extracting rows 3 to 16 of the ‘data’ field in the struct file. No sophisticated or manual artefact removal technique was employed. This was to facilitate online classification of mental workload in practice. However, eye blinks, muscle artefacts and powerline noise were removed by bandpass filtering. The dataset was passed through a low pass Butterworth filter with cut-off frequency of 3Hz. The filtering was done forward and then backward. This two-way filtering ensures zero-phase filtering or avoids phase distortion of the EEG data. Then, the low-passed data was filtered in both directions again using a high-pass Butterworth filter with cut-off frequency of 37Hz. With the aid of the markers sent during the EEG recording, the epochs corresponding to each task were extracted.

#### 2) Feature Extraction

The filtered data were divided into 4-second blocks with 2-second overlaps between adjacent blocks. The data in each block was normalised for zero mean as shown in (1) below.

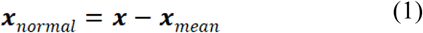

where **x** is the whole data in a 4-second block, is *x* _*mean*_ the mean of the data in such block and *x* _*normal*_ is the normalised data in the block. Normalisation gives equal importance to every data and facilitates further processing. The power spectral density (PSD) in each normalised block was computed using Welch’s method with 1-second Hamming window and 50% overlap. Windowing was necessary to obtain a short-time Fourier transform that captures the nonstationarity of EEG. In each block, the power spectral densities of eight frequency bands were computed thus: 4-8Hz, 8-12Hz, 12-16Hz, 16-20Hz, 20-24Hz, 24-28Hz, 28-32Hz, 32-36Hz. For each frequency band, the root-mean-square (RMS) value was calculated as follows:

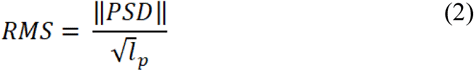

where ‖PSD‖ is the Euclidean length of the PSD in a frequency band and *l*_*p*_ is the length of the PSD vector. With 14 channels and 8 frequency bands, 112 (14 × 8) features were generated.

#### 3) Data Classification

SVM was used for the classification of the workload levels in the n-back and arithmetic tasks. The accuracy of the model was evaluated using 10-fold cross-validation. Cross-validation reflects an unbiased performance of the algorithm and prevents overfitting [35]. In addition, the performance of the SVM was investigated for cross-session, cross-task and cross-subject classifications. ASFM was also applied and the results of the two methods were compared.

ASFM was proposed in [28] as a fast domain adaptation technique for EEG-based emotion recognition to overcome the degradation of algorithm when EEG data are sampled from different subjects or sessions. The nonstationary nature of EEG and variability of brain dynamics with individuals and age causes a mismatch between the marginal and conditional distributions of the source domain (training data) and target domain (testing data). ASFM uses a framework which reduces such mismatch. In other words, if there is a source domain X_s_ with label Y_s_ and a target domain X_t_ with label Y_t_, ASFM formulates a new feature to reduce the marginal distribution mismatch between P_s_(X_s_) and P_t_(X_t_), and conditional distribution mismatch between P_s_(Y_s_|X_s_) and P_t_(Y_t_|X_t_). For details of the algorithm, see [28].

## III. RESULTS

### A. Subjective Measure (RSME)

As expected, the RSME scale revealed that mental workload perceived by subjects increased with memory load, with average rating of 44 for 0-back task and 86 for 2-back task. Paired-samples t-test showed that the two task levels were significantly different (t(7) = -9.361, p<0.05). The average ratings for the 1-digit and 3-digit arithmetic tasks were 42 and 73 respectively, and the two task levels differed significantly (t(7) = -4.47, p<0.05). The averages and statistical tests of both the n-back tasks and arithmetic tasks confirmed that the experiment design used in this work provides two discriminative workload levels: low workload and high workload.

### B. Performance Measures

Average response time increased as the workload increased from 0-back (547.9ms) to 2-back (853.9ms). Wilcoxon signed-rank test showed a significant difference in response times of the two workload levels (p<0.05). The arithmetic tasks also showed increase in response time with workload from 972.5ms (1-digit) to 1251.8ms (3-digit). The difference was statistically significant (t(7) = -4.773, p<0.05). The results imply a significant interaction between speed of performance of a task and workload.

The average accuracy of response to stimuli decreased with increase in workload from 0-back (98.2%) to 2-back (91.4%). The accuracy varied significantly between the two workload levels (t(7) =3.399, p<0.05). Increase in workload from 1-digit to 3-digit arithmetic resulted in decrease in average accuracy from 92.5% to 78.5%. Wilcoxon signed-rank test showed that the accuracy of task performance differed significantly for the two work load levels (p<0.05). The results from both n-back tasks and arithmetic tasks confirmed the expected difference between the difficulty levels of low and high mental workloads.

### C. Variation of EEG Spectral Power with Workload

The grand averages of spectral powers across all the eight subjects are shown in Fig. 4. The channels (electrodes) presented in the figures were chosen in such a way that all the brain regions are represented.

**Fig. 4.**
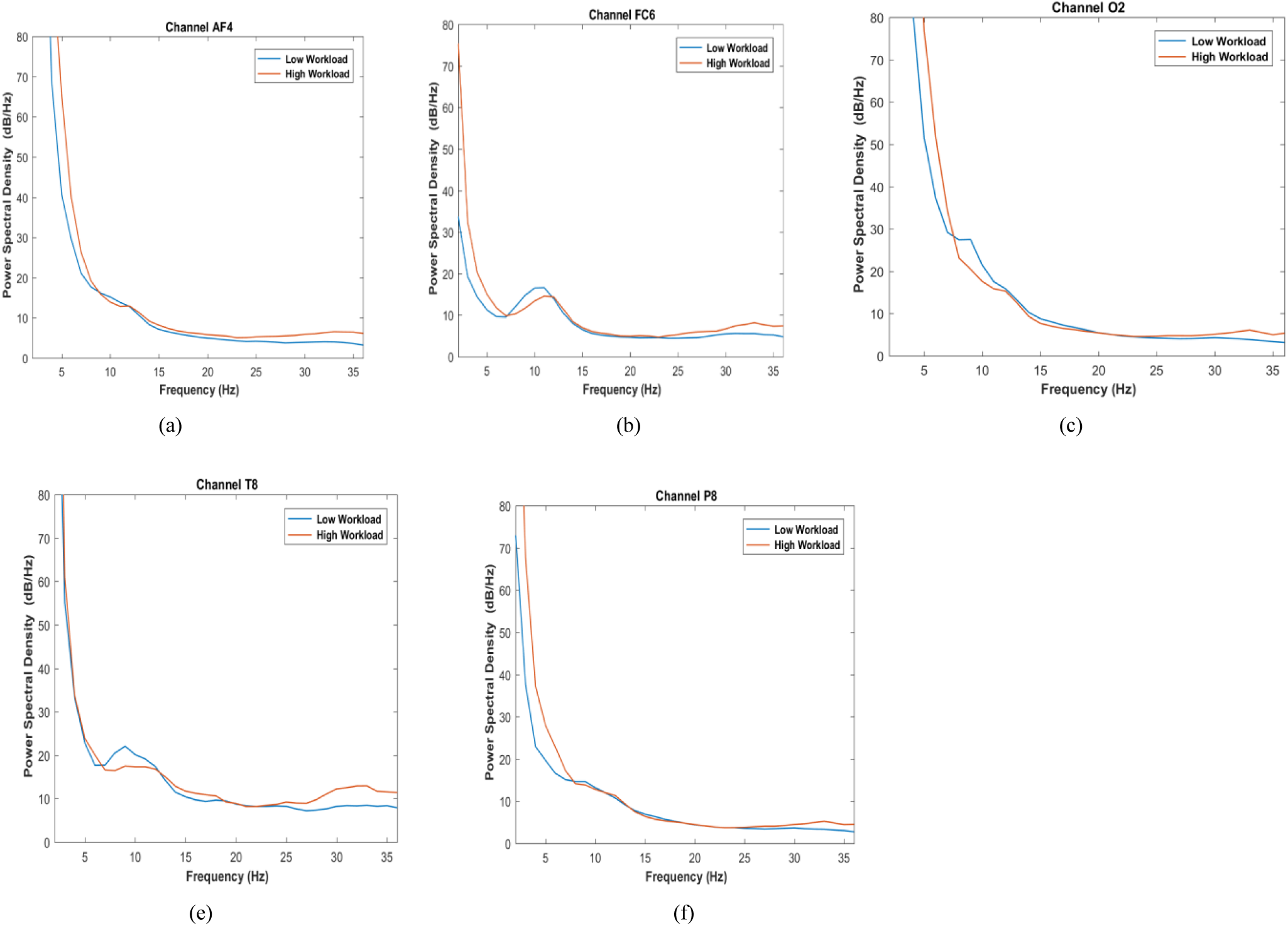
Grand averages of spectral power for the eight subjects vary with workload across frequency bands (a) Power spectra in frontal region. (b) Power spectra in central region. (c) Power spectra in Ocipital region. (d) Power spectra in temporal region. (e) Power specta in parietal region

All the plots show that EEG signal power varied with workload. In consonance with previous studies, alpha power (4-8Hz) was observed to decrease with workload across all the electrodes, gamma power (>25Hz) increased with workload, and theta power (4-7Hz) increased with workload. Similar to the findings in [8], increase in power with workload could also be observed in the high beta band (20-25Hz), especially at the frontal sites (AF4 and FC6). Furthermore, the effect of workload on spectral power is prominent in the gamma band across all electrodes. The results have demonstrated that EEG spectral power is a good feature for estimating mental workload.

### D. Classification Results

#### 1) Within-Session Classification

The EEG obtained from a subject in an experiment session was used to train and test the accuracy of the model for such subject. 10-fold cross-validation was used. Fig. 5 shows the performance of the SVM (with linear kernel) in classifying the workload levels for the two types of task.

**Fig. 5.**
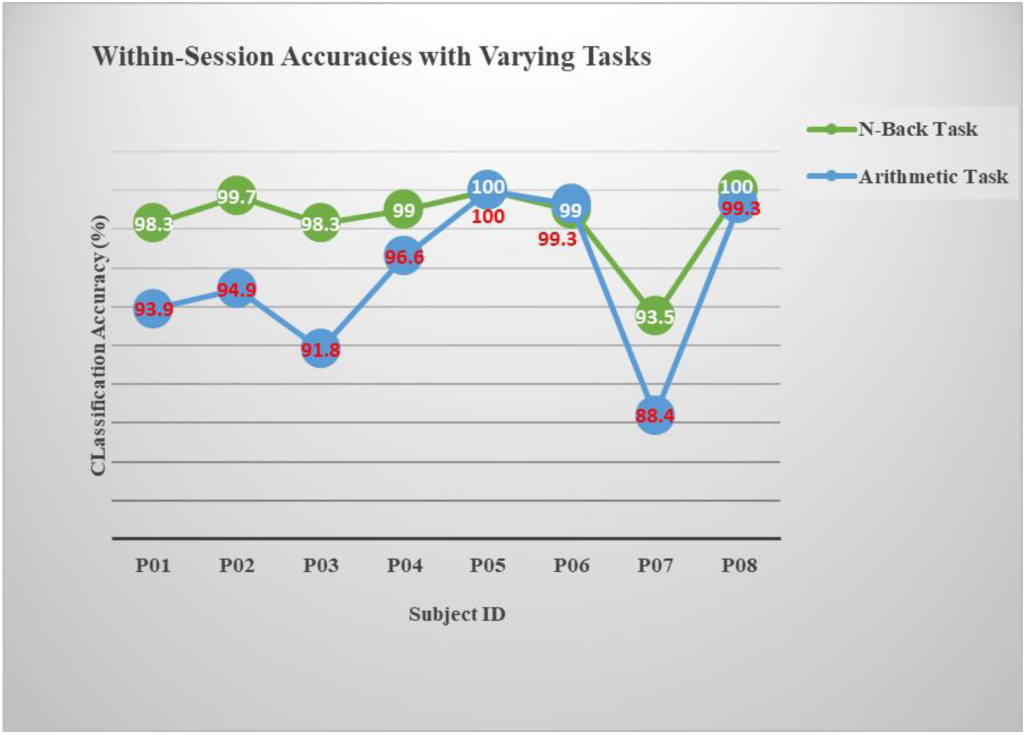
Trend of classification accuracies with tasks

The algorithm was able to classify the two levels of workload in n-back task with an average accuracy of 98.5% (SD = 2.1%) as against a mean accuracy of 95.5% (SD = 4.1%) for the two workload levels in arithmetic task. The average accuracies are close to the 98.6% (0-back vs 2-back) and 94.2% (1-digit vs 2-digit multiplication) obtained in [34] using same Emotiv EPOC headset. About 100% accuracy was also reported by [35] using Emotiv headset for 0-back vs 2-back tasks. Wilcoxon signed-rank test showed that the classification accuracies in the n-back and arithmetic tasks were significantly different p=0.028 (p<0.05). The highest and lowest accuracies achieved for the n-back task were 100% and 93.5% respectively. The arithmetic task produced 100% and 88.4% as the highest and least accuracies respectively.

The difference in accuracy for the two tasks could be attributable to the fact that the two levels of workload in the arithmetic tasks have more similarities than those of the n-back tasks. Hence, there are likely more common features in the 1-digit and 3-digit subtasks which makes less easy for the algorithm to discriminate between the two arithmetic workload levels. It could also be that there are more cross-subject variabilities in the arithmetic task than the n-back task, therefore, the model could generalise better for the latter task. As shown in the results, accuracies of the two tasks varied from subject to subject. Very high accuracies were achieved for subjects **P05** and **P08**. These discrepancies point to the fact that brain dynamics vary with individuals and age; hence, a model may not generalise well across subjects and may therefore require tuning for every user. However, the model developed in this work was good enough to generalise across many subjects without individual-based tuning.

#### 2) Classification Accuracy with Varying Data Length

To investigate the performance of the model on small number of samples, the time window of EEG signal used in cross-validation was varied from 60seconds to 300seconds. Fig. 6 shows the effect of varying data lengths (windows) on the performance of the model.

**Fig. 6.**
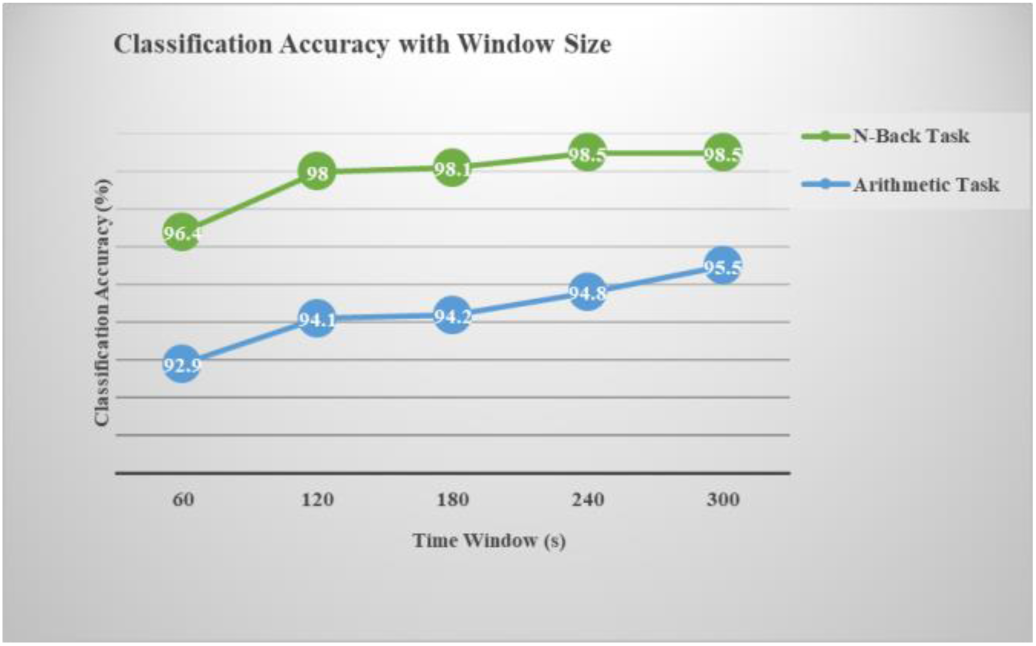
Variation of model accuracy with length of training data

As expected, the average classification increased with the amount of training data. However, the model achieved an average accuracy of 96.4% (SD = 5%) and 92.9% (SD = 7.5%) on 60second data in the n-back task and arithmetic task respectively. The result suggests that the model can achieve a high accuracy when trained with small number of data. This is very important in online usage where fast detection of workload levels is necessary. Also, if the model needs to be trained for every user at every usage session, the required time for training would not be large. Consequently, the burden of retraining model across users and sessions will not be significant.

#### 3) Cross-Session Classification

Due to the nonstationary nature of EEG signals, the performance of a model degrades if the training and test data are from different sessions or times. As a result, training is often repeated for every session. To test the performance of the model across different training sessions, three participants were asked to repeat the tasks after seven days. Then, the data from the first day were used to train while the data from the eighth day were used for testing. Here, SVM and ASFM were used and compared against each other as shown in Fig. 7 and Fig. 8.

**Fig. 7.**
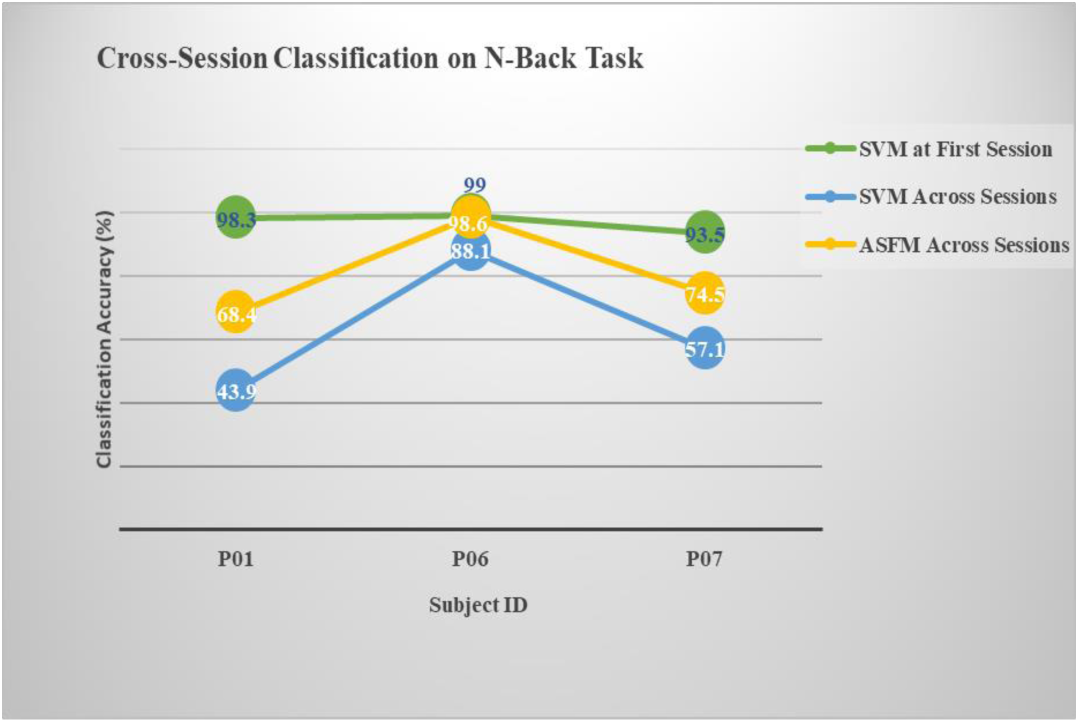
Cross-session classification on n-back task

**Fig. 8.**
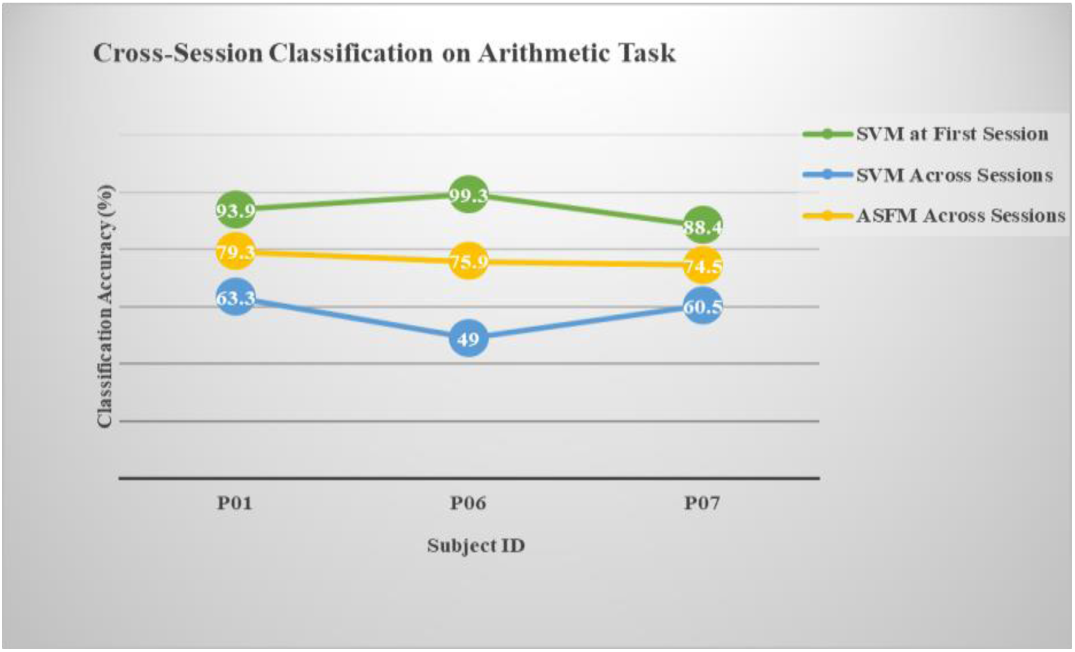
Cross-session classification on arithmetic task

The performance of SVM degraded when the trainings from previous experiment session were used to classify data obtained many days later without retraining. The accuracy of SVM, without any feature adaptation, reduced to as low as 43.9% (below 50%) in one of the cases. However, the use of ASFM, a domain adaption technique, achieved high average cross-session accuracies of 76.6% (SD = 2.5%) and 80.5% (SD=16%) in the arithmetic and n-back tasks respectively.

ASFM reduced the marginal and conditional distribution mismatch of EEG data across two different experimental sessions. This result suggests that the model with ASFM could be used for a subject at every session without retraining. ASFM was first used in [28] for emotion recognition using differential entropy as features. In that work, it achieved a cross-session accuracy of 75.1% (SD = 7.7%). This work has however shown that it can be successfully applied to mental workload using power spectral density as features.

#### 4) Cross-Subject Classification

Having known that brain dynamics vary with individual and age, the model was evaluated for cross-subject performance. The leave-one-subject-out classification method was used. Data from one subject was used for testing while the data from the remaining seven subjects were used for training. The procedure was repeated eight times so that data from every subject was used for testing. To limit the size of the training data, only about 60second data window (60 samples) was selected from each subject for inclusion in the training set. Hence, the training set contained 420 samples. In the test set, the whole 5minute length of data from a subject was used. Furthermore, the kernel of the SVM was changed to RBF kernel because the linear kernel could not find a linear hyperplane for one of the cases. As a result, SVM with RBF kernel was compared against ASFM as shown in Fig. 9 and Fig. 10.

**Fig. 9.**
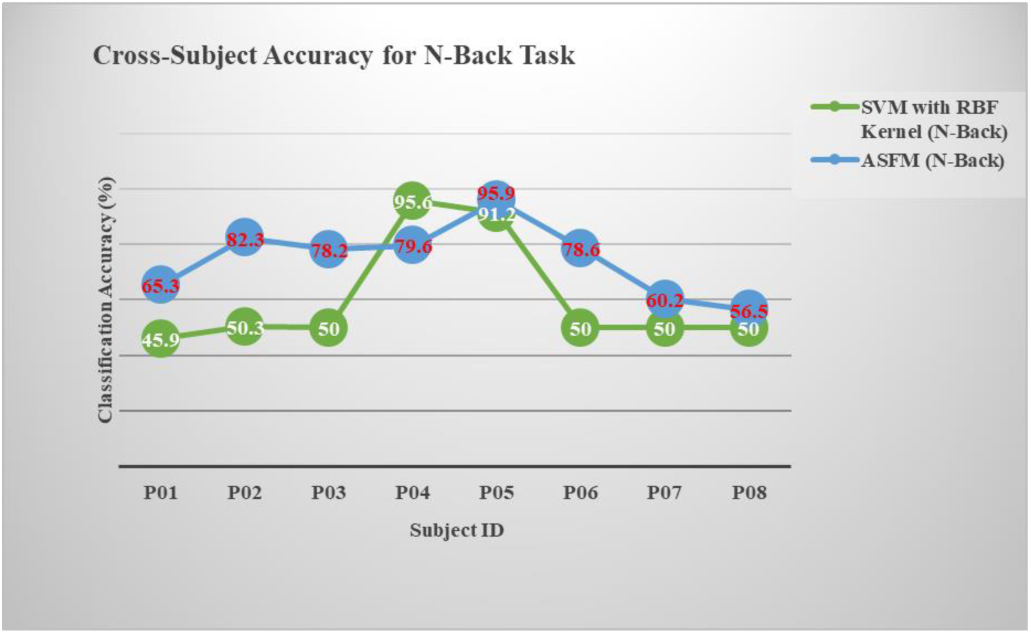
Accuracy of the model in cross-subject classification (n-back task)

**Fig. 10.**
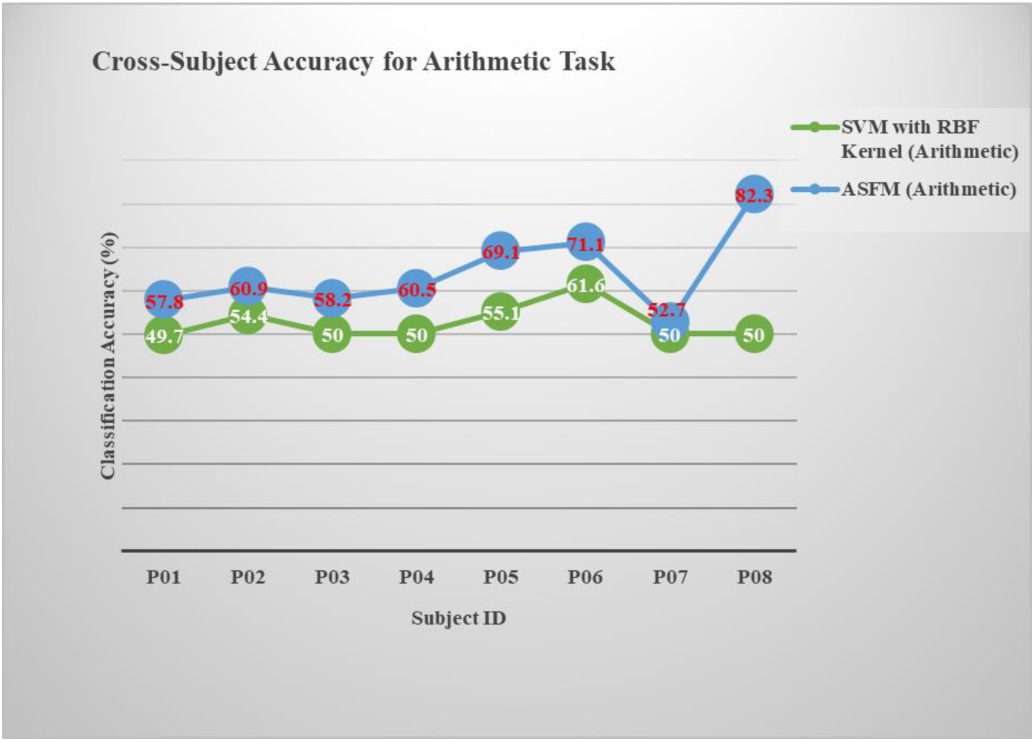
Model accuracy in cross-subject classification (arithmetic task)

SVM achieved a mean classification accuracy of 60.4% (SD = 20.5%) and 52.6% (SD = 4.2%) on n-back and arithmetic tasks respectively. ASFM improved the cross-subject accuracies to 74.4% (SD =13) and 64.1% (SD = 9.5%) in the n-back task and arithmetic task respectively. Statistical tests showed significance (or marginal significance) in the accuracies provided by the two models (p<0.05) in the n-back task and (p = 0.05) in the arithmetic task.

Even though using a non-linear kernel can improve performance of SVM or even find a solution where using linear kernel is infeasible, the performance of SVM, without feature adaptation, is limited in capturing the cross-human variability that exists in brain dynamics. Such limitation is observed in subject **P01** where the model performance deteriorated below the average level. The results have shown that feature adaptation with ASFM can mitigate the effect of subject variability on model performance.

#### 5) Cross-Task Classification

The cross-task performance of the model was examined by training on n-back tasks and classifying on arithmetic tasks. The result of the cross-task classification is shown in Fig. 11. SVM with RBF kernel provided an average accuracy of 52% (SD = 5.5%) while ASFM yielded a higher average accuracy of 68.6% (SD = 15.8%). The deterioration in performance could be attributed to difference in absolute workload levels in the two tasks. For example, low workload level in the n-back task (0-back) might not be equivalent to low workload level in the arithmetic task (1-digit). This effect can be observed in the differences of subjective ratings on the RSME scale presented earlier. Besides, the underlying brain dynamics resulting from performing the n-back tasks could be different from those of the arithmetic tasks. Nevertheless, the use of ASFM as a feature adaptation technique reduced the mismatch between the different workload types.

**Fig. 11.**
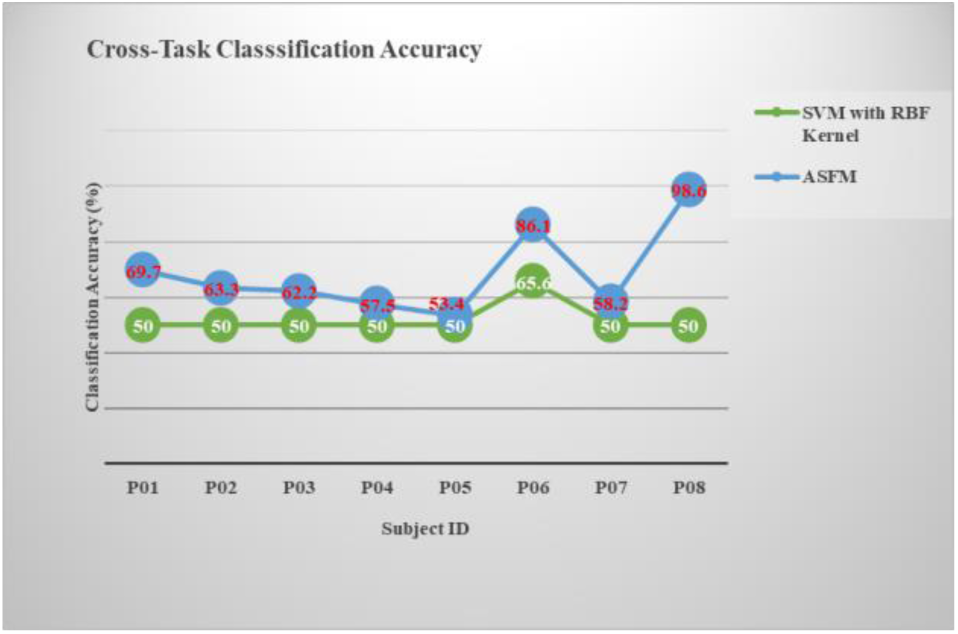
Accuracy of the model in cross-task classification

## IV. CONCLUSION

This work proposed a fast modelling technique for online estimation of mental workload using a 14-channel wireless EEG headset. The subjective and performance measures indicated that the experimental design provided discriminative workload levels. Using SVM with linear kernel, the model was able to classify workload levels in more than one type task without the need for individual or task adaptation. Furthermore, a domain adaptation technique, ASFM, was used to overcome the variabilities that exist across subjects, experimental sessions, and tasks. ASFM showed better performance than SVM (with RBF kernel) in the presence of these variabilities. ASFM – to the best of our knowledge – has not been used in estimating workload before. However, it was successfully applied in this work and yielded good performance in cross-subject, cross-session and cross-task classifications of workload. This research provided a promising framework for estimating mental workload across subjects, sessions and tasks. In addition, it shows the feasibility of developing models that would not require retraining or recalibration when there are changes in users, sessions, or types of task.

Although, this work has used two separate types of task to estimate workload, more tasks can be designed to further investigate the generalisation of the model across different tasks. In addition, multi-class workload levels can be used instead of the two-class workload levels employed in this work. This will capture more levels of workload like ‘very low’, ‘very high’, etc. Validation on many tasks and work load levels can lead to developing a task-independent model for within-task and cross-task classification in practical situations. Furthermore, other physiological markers of workload such as electrocardiogram (ECG) and electrooculogram (EOG) can be combined with EEG. Such combination can enhance exploring more features that will be robust to the variabilities of EEG. As a result, better cross-task, cross-subject and cross-session classifications could be achieved. In addition, functional connectivity model of the brain can be employed to understand the common and distinguishing features that exist in different types of tasks. Such understanding would facilitate feature selection for better cross-task classification.

Applying a model for practical situations requires validation on many samples. The model developed in this work may be tested on more subjects to validate its generalisation ability. The model can also be tested in real-time when the subjects are performing cognitive tasks. Ultimately, this can lead to a robust personalised online cognitive monitoring system for assessing mental workload in practical situations.

## ACKNOWLEDGMENT

This work was supported in part by the Petroleum Technology Development Fund (PTDF), Nigeria.

